# Is the LAN effect in morphosyntactic processing an ERP artifact?

**DOI:** 10.1101/218594

**Authors:** Sendy Caffarra, Martha Mendoza, Doug Davidson

**Author notes:** Corresponding author: Sendy Caffarra, BCBL, Basque Center on Cognition, Brain and Language, Donostia, Spain Paseo, Mikeletegi 69, 20009 Donostia-San Sebastian, Spain.

## Abstract

The left anterior negativity (LAN) is an ERP component that has been often associated with morphosyntactic processing, but recent reports have questioned whether the LAN effect, in fact, exists. The present project examined whether the LAN effect, observed in the grand average response to local agreement violations, is the result of the overlap between two different ERP effects (N400, P600) at the level of subjects (n=80), items (n=120), or trials (n=6160). By-subject, by-item, and by-trial analyses of the ERP effect between 300 and 500 ms showed a LAN for 55% of the participants, 46% of the items, and 49% of the trials. Many examples of the biphasic LAN-P600 response were observed. Mixed-linear models showed that the LAN effect size was not reduced after accounting for subject variability. The present results suggest that there are cases where the grand average LAN effect represents the brain responses of individual participants, items, and trials.

## 1 Introduction

Neural processes of sentence comprehension can happen very quickly after the presentation of linguistic input. Electrophysiological techniques, such as EEG or MEG, can monitor brain responses as they unfold over time and identify neural activity associated with cognitive mechanisms. The left anterior negativity (LAN) is a negative evoked component typically observed around 400 ms after the presentation of a target word embedded in a sentence, with a focus at left and anterior electrodes for commonly-used reference locations (i.e., mastoids). The LAN effect is an increased average negative potential which can be observed in response to a morphosyntactic violation as compared to the corresponding correct sentence (it has also been observed with increasing morphological complexity in the absence of violations; e.g, Krott & Lebib, 2013). This effect is usually followed by a greater posterior positivity (P600) and it has been reported in response to local agreement violations in morphologically rich languages (Molinaro, Barber, & Carreiras, 2011; Molinaro, Barber, Caffarra, & Carreiras, 2015; but see: Wicha, Moreno, & Kutas, 2004). The LAN effect has high theoretical relevance in some models of sentence comprehension, being related to morphosyntactic analysis (Friederici, 2002). Regardless of its functional significance, the existence of the LAN as a real ERP effect has been recently questioned^1^. Multiple ERP studies have highlighted the lack of correspondence between the LAN effect obtained after averaging the ERP responses across items and subjects (i.e., grand average) and the patterns that have been observed on a single-subject basis (Osterhout, McLaughlin, Kim, Greenwald, & Inoue, 2004; Tanner & van Hell, 2014). These findings have suggested that the LAN might be a spurious effect due to averaging procedures. Given the fact that the LAN is relevant in the sentence comprehension literature, and that averaging over subjects and items is a standard procedure in the ERP analysis, it becomes important to model the variability of this ERP effect.

The present study investigates the variability of the LAN effect that has been observed in the grand average waveforms in response to local agreement violations (determiner-noun gender agreement) in a morphologically rich language (Spanish). Three different claims supporting the idea that the LAN effect is an artifact will be explored in a large ERP dataset (80 subjects; 120 items per condition).

The first claim states that individual subjects and individual items do not consistently show a LAN effect, although this effect is evident in the grand average waveforms (Osterhout, 1997; Osterhout et al., 2004). According to this view, around 400 ms after stimulus onset the individual ERPs show a continuum between two distinct ERP effects, neither of which are LAN effects, but rather the N400 and the P600. The N400 is a negative component that (also) peaks around 400 ms after stimulus onset but has a focus over central-posterior electrodes (Kutas & Federmeier, 2011), rather than left anterior electrodes. As we have described, the P600 is a posterior positivity (which can be right lateralized; Tanner & Van Hell, 2014) that is usually observed 500 ms after stimulus onset, although there is certain variability in the timing of its onset (Osterhout & Mobley, 1995), with some subjects showing an earlier onset, and some a later onset. Specifically, the first claim is that during averaging over subjects, the spatial and temporal overlap of these two qualitatively different effects would reduce the original N400 effect over (right-) posterior sites, producing a residual (left-) anterior negative effect in the grand average (Osterhout, 1997; Osterhout et al., 2004; Tanner & Van Hell, 2014) ^2^. This claim will be tested by quantifying the LAN, the N400 and the P600-like effects at the subject as well as the item level between 300 and 500 ms. Since the same argument can be applied to single trials within the same participant, similar analyses will be carried out over small sets of trials.

The second claim is that there is a low co-occurrence of the LAN and the P600 effects across individuals (Osterhout et al., 2004; Tanner & Van Hell, 2014). Single subjects would typically show negative-dominant or positive-dominant responses, rather than biphasic responses. According to this view, the sub-population of participants with large negative effects tend to show small P600 effects, while the sub-population having a clear P600 effect would show small early negativities. Hence, the co-occurrence of negative and positive effects would be present in a small subset of participants and it would mainly arise from averaging over participants. This claim will be tested by calculating the correlation between early dominant negative effects and later posterior positive effects across subjects. Since the same argument can be applied to single items, similar analyses will be carried out across items.

The third claim is related to those described above and it frames the idea that the LAN is the result of subject variability in general. Note this is independent of whether there is a biphasic LAN-P600 response or rather monophasic LAN and P600 responses. The logic is the following: If the effect observed in the grand average is driven, at least in part, by individual differences (Osterhout et al., 2004; Tanner & Van Hell, 2014) then a model that includes specific terms for subject variability should provide a better fit than a model without those terms, subject to a penalty for the additional terms. Also, a model that accounts for subject variability should account (at least partially) for the LAN effect size, which is defined as the fixed effect coefficient over its SE (Gelman & Carlin, 2014). Hence, a reduction of the LAN effect size should happen through a reduction of the fixed effect estimate and/or through an increase of the relative SE. This claim will be tested by fitting linear mixed effects models on the LAN voltage and comparing them. Similar models will be also fit for the P600 voltage in order to see whether individual differences specifically account for the LAN effect.

### 1.1 The present study

The above-mentioned claims were tested using a multi-level modeling approach that characterizes the statistical properties of both subject and item level. We used an ERP dataset from three previous experiments focused on the LAN effect (Caffarra & Barber, 2015; Caffarra, Barber, Molinaro, Carreiras, 2017). The experimental materials and designs in these studies were always the same and included gender agreement violations between a prenominal determiner and a target noun, a morphosyntactic violation that often elicits a LAN effect in the grand average of highly-proficient subjects (Molinaro et al., 2011; but see: Wicha, Moreno, & Kutas, 2004). The ERPs time-locked to the target noun were calculated across each item, each participant, subset of trials, and overall (for methodological details see below). The ERP effects were defined as the difference between the disagreement and the agreement condition.

According to the first claim, between 300 and 500 ms after the target onset there should be a discrepancy between the ERP effect observed in the grand average and the ERP effects of single subjects (or single items, or small sets of trials). The topographies of by-subject, by-item, and by-trial effects should mainly show central-posterior negative effects (N400) and posterior positivities (P600). The LAN effects were not expected to be elicited by the majority of participants (or items, or trials).

According to the second claim, most of the participants (or items) should show either a positive or a negative dominant response, with a minority showing some degree of biphasic response (between 5% and 35% according to Tanner, 2015; Tanner & Van Hell 2014). A strong correlation should be observed between the dominant negative effects seen between 300 and 500 ms at the individual level and the late positive effects (500-800 ms), so that in subjects where one effect is larger, the other effect should be smaller (0.59<|r|<0.74; in case the dominant effects seen at the individual level have the same posterior location; Tanner, Inoue & Osterhout, 2014; Tanner, McLaughlin, Herschensohn, & Osterhout, 2013; Tanner & Van Hell, 2014).

The third claim predicts that LAN effect size estimate should be reduced and/or the SE should increase after including in the model a term for subject-specific violation effects. Adding this random effect should also lead to an improvement of the model fit, as this would provide evidence for some subject-to-subject variability in the LAN effect.

## 2 Material and Methods

Eighty Spanish speakers highly-proficient in Spanish participated to the studies (45 female; mean age=25 y; mean Spanish AoA=1.4 y; average proficiency score=4.9/5.0). They read 240 Spanish sentences that were presented word by word in rapid serial presentation. Half of the sentences contained article-noun gender agreement violations and half were grammatically correct. After each trial they performed a grammaticality judgement task (mean accuracy=95.4%; SD=3.0). Further details about the participants, the procedure, the EEG recording and preprocessing can be found in the original papers (Caffarra & Barber, 2015; Caffarra et al., 2017; for the EEG preprocessing we followed the pipeline described in Caffarra & Barber, 2015^3^ and data from Caffarra et al., 2017 were down-sampled to 250 Hz). ERPs from two experimental conditions (agreement and disagreement) were considered. Overall, 8.7% (SD=6.1) of trials were excluded due to artifacts or incorrect behavioral responses. The number of remaining trials did not differ between conditions (*t*(79)=1.3, *p*>.05).

Grand average waveforms, by-subject, and by-item ERPs were calculated for each condition. For the first analysis, the topographies of the ERP effects (i.e., disagreement-agreement voltage difference) between 300 and 500 ms were obtained at the subject and item level. In order to reduce the potential artifacts generated by averaging procedure, multiple ERP effects were also calculated on the smallest possible trials sets (size range: 1-2 trials) for each participant (total number of sets: 6160, see Supplementary Materials A).

The number of subjects, items and trial sets showing a LAN, an N400, or a positive effect was calculated. The presence of a LAN effect was defined as a negative effect over a representative left-anterior site (F3), which was greater than the effect registered over a posterior site (Pz). The presence of a N400 effect was defined as a negative effect over a posterior site (Pz) being greater than the effect over a left-anterior site (F3). Any other case where both F3 and Pz showed a positive ERP difference was categorized as a positive effect^4^. As an aside, note that the distinction between LAN and N400 effect is based on topographical distribution features (Kemmerer, 2015), which are not straightforwardly related to neural sources in the absence of an adequate forward model (Hallez et al., 2007). Hence, our results should not be read in terms of ERP source modelling.

For the second analyses, the Pearson correlations were calculated between the LAN voltage difference (disagreement-agreement ERP difference at F3 between 300 ms and 500 ms) and the P600 voltage difference (disagreement-agreement ERP difference at Pz between 500 and 800 ms) across subjects and items. Similar correlations were calculated between the N400 voltage difference (disagreement-agreement ERP difference at Pz between 300 ms and 500 ms) and the P600 voltage difference (as in Tanner et al. 2013, 2014; Tanner & Van Hell, 2014).

For the third analysis, mixed-linear models (Baayen, Davidson & Bates, 2008) were fit using the LAN-time-window voltage (ERP average amplitude at F3 between 300 ms and 500 ms) as the dependent variable and agreement as a fixed effect factor at the level of individual trials. By-item and by-subject random intercepts, as well as random slopes for agreement were incrementally added to model variability in the LAN responses at the level of individual subjects and individual items, in addition to the variability of individual trials already modeled by the fixed effect terms. The contribution of each random effect to the model fit was assessed using likelihood ratio statistics (Pinheiro & Bates, 2000; Bates et al. 2014) by comparing the respective model with one that was identical except for the effect in question. Similar analyses were carried out using the P600 voltage (ERP average amplitude at Pz between 500 ms and 800 ms) as a dependent variable. A final model on the LAN voltage was also performed including the subject-specific N400 violation effect term. Dummy coding was used to label the levels of the categorical predictor (0=agreement; 1=disagreement). Analyses were carried out using the *lme4* package (Bates, Maechler, Bolker & Walker, 2014). *P*-values for the coefficients of the linear mixed-effect models were calculated treating the *t*-statistic as if it were a z-statistic^5^. Note that inferences about model fit were not based exclusively on likelihood ratio statistics, *p*-values or confidence intervals, but also on a variety of other factors (Loken & Gelman, 2017).

## 3 Results

The grand average waveforms showed a clear LAN-P600 biphasic response (Figure 1; Caffarra & Barber, 2015; Caffarra et al., 2017).

**Figure 1.**
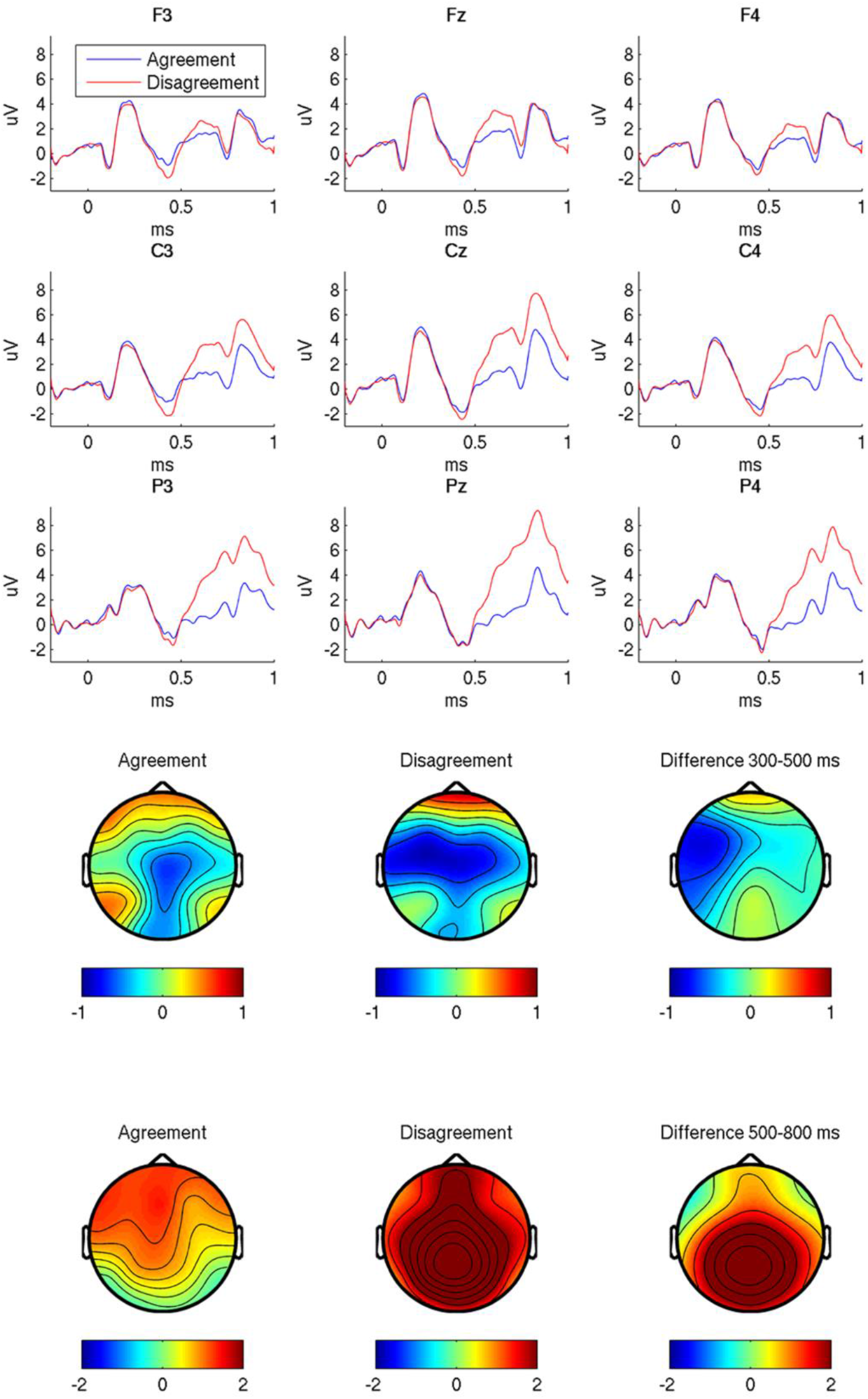
The upper panel shows grand average waveforms for the agreement and the disagreement condition (blue and red line, respectively). Positive voltage is plotted upwards. The bottom panel shows the topographic distribution of each experimental condition and of their difference in the two time windows of interest: 300-500 ms and 500-800 ms. Y-axes units are microvolts, x-axes are ms, and the topography color-scales are microvolts. The greatest magnitude LAN effect is at F3, and the greatest P600 effect is at Pz.

The analysis of the subject topographies revealed that 55% of the participants showed a LAN effect, 25% a N400 effect and 20% a positive effect (Figure 2). The by-item topographies showed that 46% of the items elicited a LAN, 29% an N400 effect and 25% a positive effect. Similar percentages were observed after calculating ERP effects on small sets of trials for each participant (LAN: 49%, N400: 29%, positive effect: 22%, see Supplementary Materials A). Negative effects at F3 were present in 71% of the participants and 66% of the items (Figure 2).

**Figure 2.**
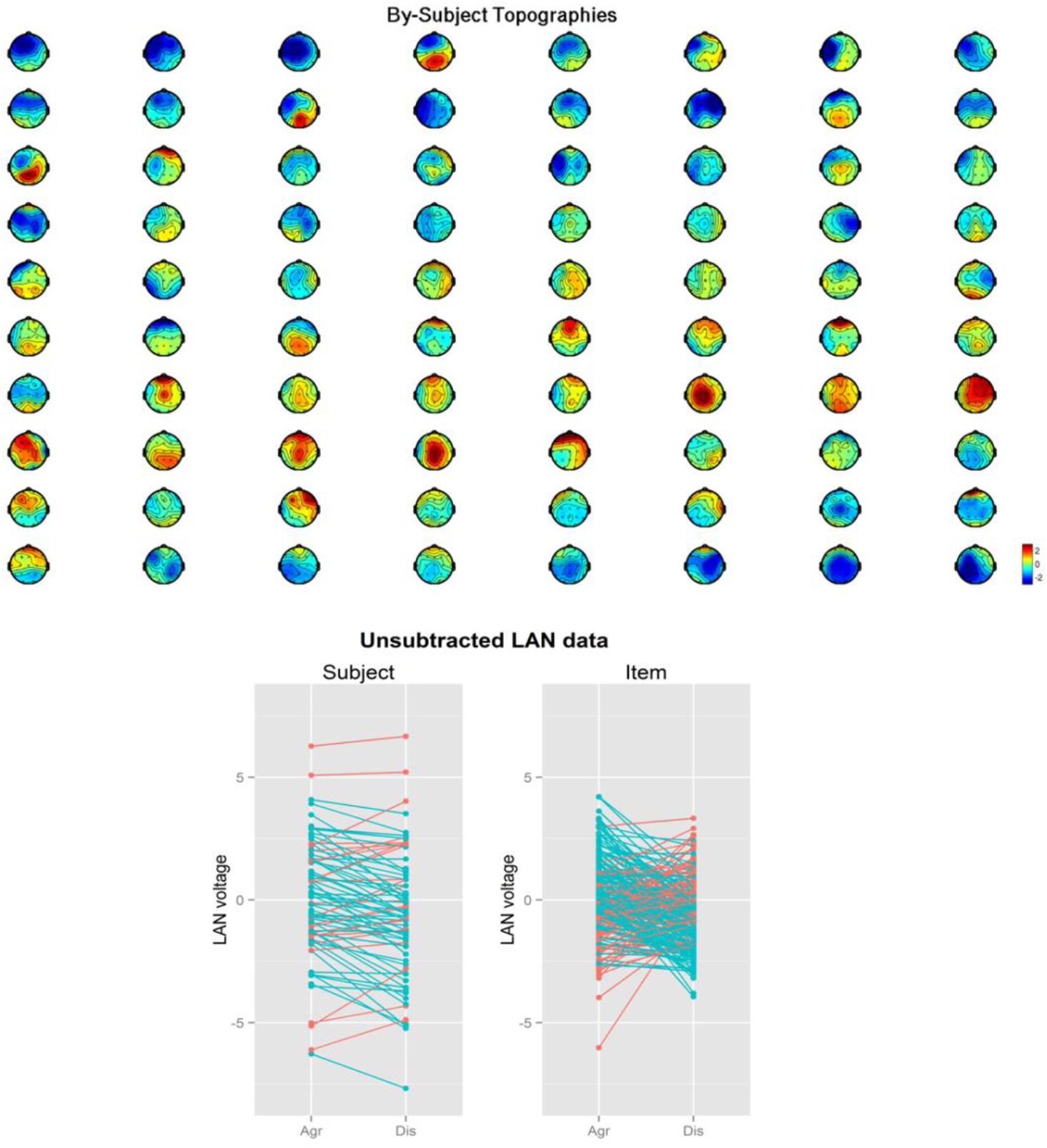
The upper panel shows the topographic distributions of the ERP effects between 300 ms and 500 ms for each participant. The topographies are ordered to show the LAN effects (44, from the greatest effect to the smallest one), the positive effects (20) and the N400 effects (16, from the smallest effect to the greatest one), respectively. The bottom panel shows the ERP voltage recorded at F3 for the agreement and the disagreement condition. Blue lines marked a negative ERP effect, while red lines marked a positive effect.

In the second analysis, 66% of the participants and 62% of the items showed an early negative effect at F3 followed by a later positivity (Figure 3, first row, quadrant IV corresponds to the presence of biphasic responses). The LAN effect and the P600 effect were uncorrelated across participants (*r*=0.04; *p*=0.72). However, there was a positive weak correlation across items (*r*=0.25; *p*<0.001), suggesting that for items where the P600 effect was smaller, the LAN effect was larger. When similar correlations were computed on the ERP voltage differences at the same representative electrode (Pz) larger correlations were found (by-subject: *r*=0.48;*p*<0.001; by-item: *r*=0.64; *p*<0.001).

**Figure 3.**
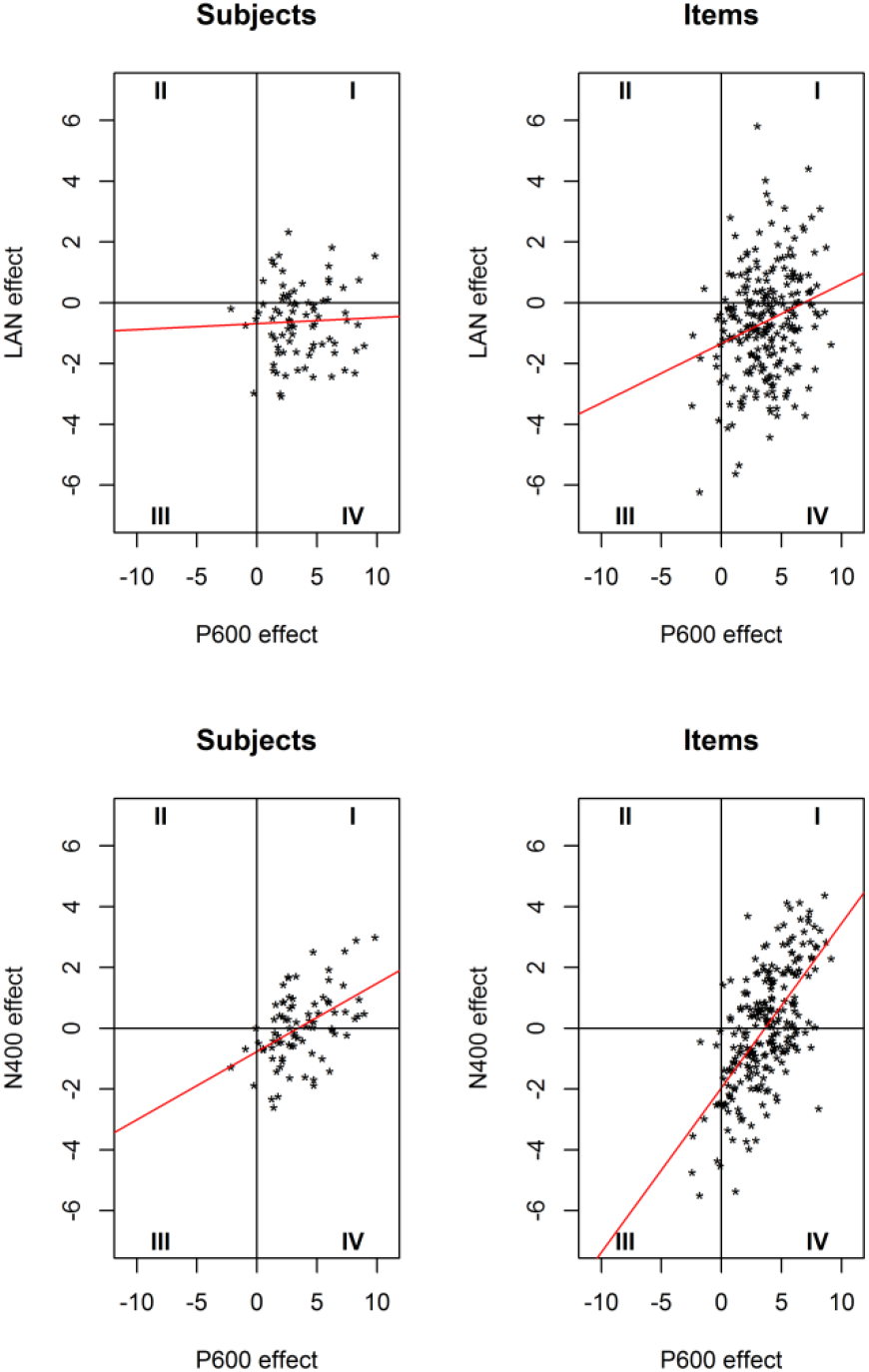
Top row: by-subjects and by-item correlations between the LAN voltage difference (disagreement-agreement ERP difference at F3 between 300 ms and 500 ms) and the P600 voltage difference (disagreement-agreement ERP difference at Pz between 500 and 800 ms). Bottom row: by-subjects and by-item correlations between the N400 voltage difference (disagreement-agreement ERP difference at Pz between 300 ms and 500 ms) and the P600 voltage difference (disagreement-agreement ERP difference at Pz between 500 and 800 ms). Each quadrant is displayed with its relative number.

In the third analysis, an initial mixed-linear model was fit on the LAN voltage including agreement as a fixed predictor and a by-item random intercept (model A: intercept=0.04; SE=0.10; β=-0.62; SE=0.13; *t*=-4.94, *p*<0.001, for more details see Supplementary Materials C). Note that the agreement condition average voltage is near zero (0.04) and that the average LAN (disagreement) effect is a negative modulation of about half a microvolt (−0.62). The standard error of the LAN effect (0.13) is small relative to the size of the effect, leading to a relatively large t-ratio. Compared to this baseline model, adding additional random effect terms would be expected to affect the standard error of the fixed effects, and possibly the fixed effect coefficients.

Adding a by-subject random intercept (model B: intercept=0.05; SE=0.28; β=-0.62; SE=0.12; *t*=-5.13, *p*<0.001) led to an improvement of the model fit (χ^2^(1)=1167, *p*<0.001) and an increase of the SE (from 0.10 to 0.28). This is consistent with larger subject variability in the LAN-window voltage in the control condition (Figure 4). To assess whether a subject-specific and/or an item-specific violation effect improved the fit, two additional models were compared: one in which a by-subject random slope was added to model B (model C: intercept=0.05; SE=0.27; β=-0.62; SE=0.13; *t*=-4.61, *p*<0.001), and one that also included a by-item random slope (model D: intercept=0.05; SE=0.27; β=-0.62; SE=0.14; *t*=-4.57, *p*<0.001). Note first that the fixed parameters and the SEs remain unchanged. Second, neither of these models led to improvements in fit over the corresponding nested comparison (model C: χ^2^(2)=2.96, *p*=0.23; model D: χ^2^(2)=2.44, *p*=0.30). These model comparisons, along with plots of the distributions of the model random effects, as well as plots of the raw subject averages, suggest that the LAN effect was not subject to a great deal of subject-to-subject or item-to-item variability, although variability in the effect was clearly observed. Importantly, the estimates of the LAN effect (−0.62 μV, SE=0.13 μV) did not change substantially after accounting for subject variability. Finally, we also fit a model that included the amplitude of the N400 effect (at Pz) as a predictor. In this case, the estimate of the LAN effect was not reduced and the SE remained unchanged (−0.65 μV, model E: intercept=0.32; SE=0.26; β=-0.65; SE=0.13; *t*=-4.97, *p*<0.001).

**Figure 4.**
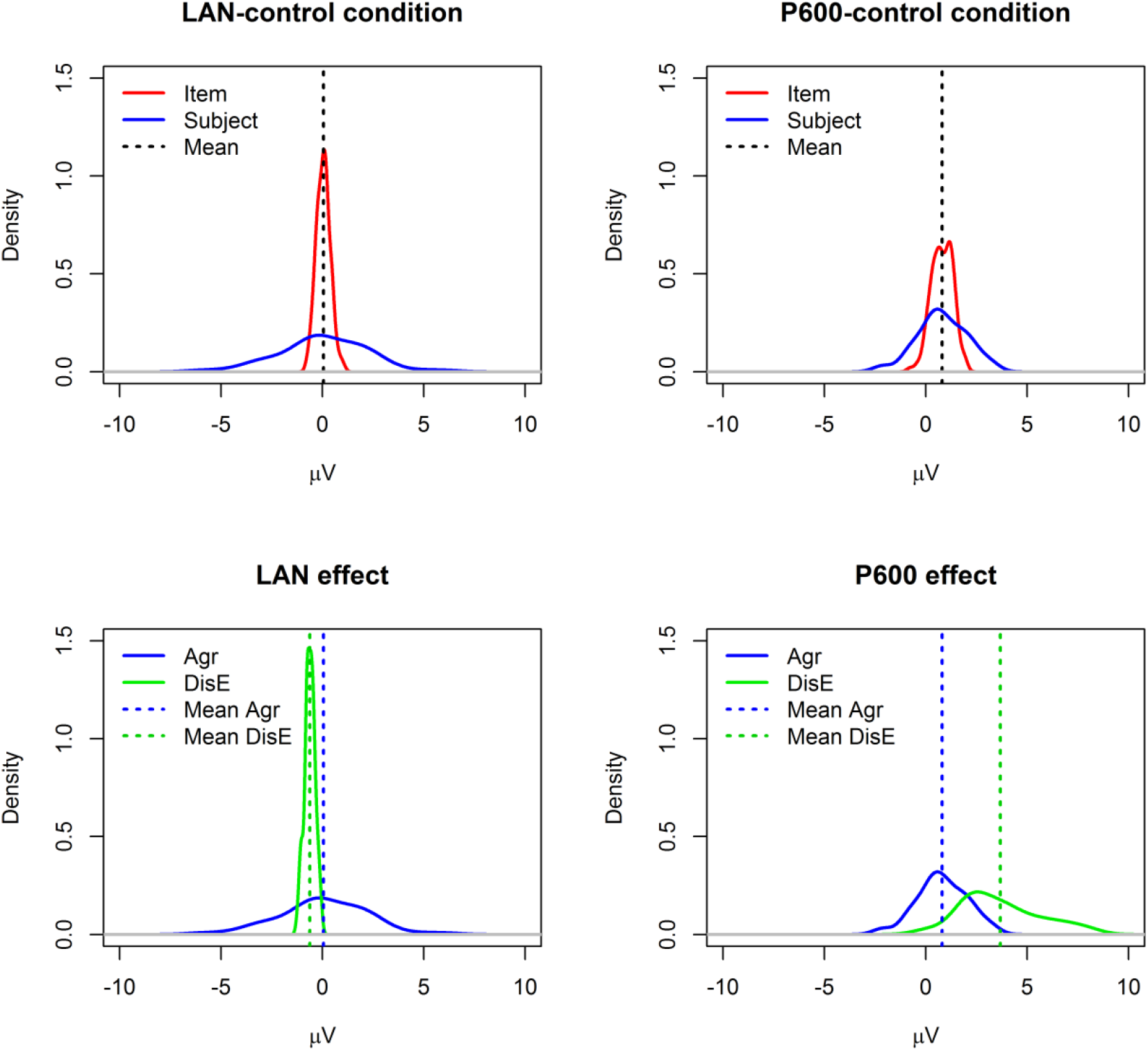
Density plots of the variance parameters estimated from models C. Y-axis are kernel density estimates, and x-axis units are microvolts. The curves represent the estimates of the population probability distribution based on the given sample of points. The upper panel shows the LAN and the P600 average response to the control condition (vertical dashed line), and variability due to subject (blue distribution) and item (red distribution). The lower panel shows the ERP average response to the control condition (Mean Agr: blue dashed line) and the ERP average size of the violation effect (Mean DisE: green dashed line) with the relative variability due to subjects. For a comparison between the distribution of the observed data and the distribution of the fitted random effect coefficients see Supplementary Materials D.

The models A, B, C, and D were also fit to the P600 voltage (500-800 ms time window; for more details see Supplementary Materials C). All models had substantially equivalent estimates of the baseline voltage (about 0.8 μV) as well as P600 violation effect (about 3.7 μV; model A: intercept=0.81; SE=0.11; β=3.69; SE=0.14; *t*=26.89, *p*<0.001; model B: intercept=0.79; SE=0.25; β=3.70; SE=0.13; *t*=27.59, *p*<0.001; model C: intercept=0.81; SE=0.19; β=3.68; SE=0.28; *t*=13.20, *p*<0.001; model D^6^: intercept=0.81; SE=0.18; β=3.67; SE=0.29; *t*=12.71, *p*<0.001).

Similar to the LAN, adding a by-subject random intercept improved the fit over model A (χ^2^(1)=629.55, *p*<0.001), with an increase of the SE (from 0.11 to 0.25). In addition, including a by-subject random slope led to an improvement of the fit over model B (χ^2^(2)=182.72, *p*<0.001) and a concomitant increase of the SE (from 0.13 to 0.28). Thus, there is evidence of subject-to-subject variability in the P600 control condition, as well as in the P600 violation effect (Figure 4).

## 4 Discussion

The present study examined and modeled the variability of the LAN effect elicited by local agreement violations in a morphologically rich language. Three different hypotheses related to the claim that the LAN effect in the grand average is a spurious effect were examined. A large ERP dataset showing a LAN-P600 biphasic response after grand averaging was analyzed at the subject, item, and trial level.

First, individual ERPs were calculated by-item, by-subject and by-trial sets to examine how often the ERP responses resembled the grand average for (a) single subjects (over all items), (b) single items (over all subjects), and (c) small sets of trials. This analysis clearly showed that LAN effects could be observed in individual subjects, items, and trial sets. Moreover, these effects were more consistent than the N400 or positive effects at the level of subjects, items and small sets of trials. The grand average LAN effect was, thus, more representative of what occurred to most subjects, items, and trials than the proposed alternatives. Note that even when the effects of averaging were kept to the minimum (in the by-trial analysis ERP averages were either not computed or computed over two trials) the LAN was still the most consistent ERP effect. This is difficult to reconcile with the claim that the LAN effect depends on the averaging procedure.

These findings do not support the claim that the averaging procedure is artificially producing a left anterior negativity (Osterhout, 1997; Osterhout et al., 2004), at least when the LAN effect is elicited by local agreement violations in a morphologically rich language with highly-proficient readers. A remaining possibility is that there is a difference between violation types, languages, and participant language background. Note that although the LAN was the most consistently reported effect, it was not present in all subjects, items, and trials. Electrophysiological effects (even well-established ones, such as N1 or P2) do not likely appear at every by-subject and by-item average (Cooper, Osselton, & Shaw, 1969; Sörnmo & Laguna, 2005). The relative size of the effect together with the amount of measurement variability and measurement noise^7^ can make the effect difficult to detect at every observation. Crucially, although the LAN has a relatively small size (around 0.5 μV) as compared to the N400 and the P600 effects (which are typically over 1 μV), it was consistently detected. As to the impact of measurement variations, the first analysis cannot tell us whether subject (or item) variability partially accounts for this ERP patterns. This was specifically tested by the third analysis.

Second, the correlations between LAN (which was the dominant negative effect based on the results of the previous analysis) and P600 single effects were calculated to examine how the biphasic responses covary over the sample of subjects or items. The results revealed that most of the participants and most of the items show both early negative and late positive effects. There was no clear relation between individual subject LAN and P600 effects, but there was a weak correlation across items. As the by-item LAN effect increased the by-item P600 effect decreased. When similar correlations were calculated on the same posterior spatial location they were stronger and compatible with those reported in previous studies (Tanner et al., 2013, 2014; Tanner & Van Hell, 2014).

These findings speak against the presence of prominent monophasic responses at the subject level (Osterhout et al., 2004), as well as at the item level. These results show weak support for the covariance of the two components of the biphasic responses. The LAN effects (which were the most frequent ERP negative effects observed in individual subjects and items) did not show a strong relation with the subsequent positive ERP effects, while ERP effects of individual items showed only a weak relation. Note that the correlations calculated here regarded different topographic locations (F3 and Pz), as the first analysis showed that early negative effects were most frequently left anteriorly distributed. However, when the same posterior topographic location was considered (as done in previous studies: Tanner et al., 2013, 2014; Tanner & Van Hell, 2014), the correlations were stronger, suggesting that the covariance of two subsequent ERP magnitude differences increases with spatial contiguity.

These higher correlations at the same topographic locations are probably due to a shared random effect and measurement noise at the same electrode.

Third, mixed-linear models were fit to estimate ERP effects after accounting for subject and item variability. The results showed that larger subject variability was present in the LAN responses to correct sentences, as well as in the P600 responses to correct and incorrect sentences. The estimate of the LAN fixed effect parameter was not reduced and the SE remained unchanged after accounting for subject-specific LAN effect and for subject-specific N400 effect.

These findings provide evidence of subject-to-subject variability in the LAN responses. However, this is not a peculiarity of the LAN and it can also be observed in the P600 responses (even to a larger extent). These results do not strongly support the idea that the subject variability accounts for the LAN effect reported in the grand average^8^. In addition, the distributions of the random effects relative to the LAN control condition and violation effect (see Supplementary Materials C) do not cluster into distinct groups that would be associated with qualitatively different effects (i.e., N400 and P600), but they rather appear to be unimodal distributions.

Overall, these results argue against the possibility that the LAN effect is consistently and systematically an artifact of averaging. Note that this is true regardless of how the LAN effect is functionally interpreted. The present findings cannot tease apart different functional interpretations of the LAN effect (e.g., morphosyntactic analysis vs. working memory demands). Instead, the present work provides evidence that there are cases where the grand average LAN effect is representative of by-subject, by-item and by-trial responses. The type of violation considered here (i.e., determiner-noun gender agreement violation in a morphologically rich language) has been often associated with LAN effects in the published literature^9^ on first language comprehension (Molinaro et al., 2011; but see: Hagoort, 2003; Wicha, Moreno, & Kutas, 2004). It is still not clear whether the present results can be generalized to other experimental designs where the grand average LAN effect seems to be less representative of single ERP responses (e.g., subject-verb agreement violations in morphologically poor languages; Tanner & Van Hell, 2014; violations where there is a longer time interval between agreeing constituents; Roll, Gosselke, Lindgren, & Horne, 2013).

A potential contribution of the present work is providing useful methodological tools to examine both the properties of the ERP grand average and individual responses. The updated scripts for the present analyses are available online at https://github.com/dvdsn/218594. Estimation of different sources of variability and larger sample sizes will help us to have a better measure of ERP effect sizes and its systematic variation (see a recent debate on sample size in Friston 2012; Lindquist, Caffo, & Crainiceanu, 2013; and also the discussion in Loken & Gelman, 2017). Note however, that there remains a great deal of work to be done, as even the best-fitting models reported here did not account for very much variability (in the sense of R^2).

To conclude, the present study described multiple ways of exploring the relation between grand average waveforms and ERPs of single subjects, items, and trials. It provided evidence that the grand average LAN effect can accurately reflect individual brain responses.

## Authors Notes

This work was supported by the Spanish Ministry [PSI 2014-54500-P; IJCI-2016-27702; PSI2017-82941-P]; the Basque Government [PI_2015_1_25]; and the Severo Ochoa [SEV-2015-0490].

## Footnotes

1 This is not the first debate about the functional nature of ERP components associated to sentence comprehension. In the past years, at least another similar debate has been proposed, which was focused on the nature of the P600 and the possibility to reduce it to a P300 (Coulson et al., 1998; Osterhout & Hagoort, 1999).

2 It is worth noting that the coexistence of the LAN with a positivity does not necessarily inform on potential component overlaps. In fact, a negative effect on the scalp is always accompanied by a positive counterpart (although not always within the sensor array) since the sum of potentials around the head should be zero in ideal conditions.

3 Using the ERP pipeline described in Caffarra et al. (2017) led to similar results (Mendoza, 2017).

4 Other ways of categorizing the effects (using clusters of three electrodes instead of one representative electrode and calculating laterality and anteriority indexes based on 26 electrodes) led to similar pattern of results (i.e., presence of LAN effects at the subject and item level, greater proportion of LAN effects than any other alternatives, see Supplementary Materials B).

5 Correlations and mixed-linear models were calculated also using clusters of electrodes instead of a single representative electrodes (LAN: F3, F7, FC5; N400/P600: Pz, P3, P4) and the results were similar to those presented here. For the mixed-linear models, the results of the model comparisons were the same after fitting the same models using sum coding (−0.5/0.5). Model comparisons with leave-one-out cross validation after fitting the equivalent Stan models using the *rstanarm* package (available online at https://CRAN.R-project.org/package=rstanarm) resulted in similar conclusions.

6 The model did not converge, but coefficient estimates were similar to the models that did converge.

7 Measurement noise refers to a pattern that would not repeat with re-test, while measurement variability refers to a pattern that does repeat with re-test, but is different from the population central tendency.

8 The results from the mixed-linear models might seem to be at odd with the scatter plots of correlation analyses, where more variability seems to be present in the case of the LAN than the P600 effect. However, it should be kept in mind that the scatterplots do not represent the same type of information conveyed by the mixed-linear results. In the mixed-linear models not all variability has been modeled, but only that which is related to item and subject identity (both residuals and *R*^2^ coefficients of mixed-linear models showed that there is still a considerable portion of unmodeled activity). In addition, unlike the scatterplots, mixed-linear models model the different source of variability for LAN and P600 voltages for each condition (and not for their difference). Finally it is worth noting that the magnitude of these two effects is different (3 μV against 0.5 μV), with the larger effect being more easily modeled against the measurement variability and noise.

9 We do not know the number of studies that have looked for the LAN but have not found it --not least because of the publication bias for statistically-significant results (see Caffarra et al., 2015 for similar concerns). Even in the case the publication bias contributes overestimating the LAN effects reported in the literature, the present dataset still represents a case where the LAN effect observed in the grand average can be observed also in individual brain responses. Future studies are needed to check how far the present findings can be generalized.

---

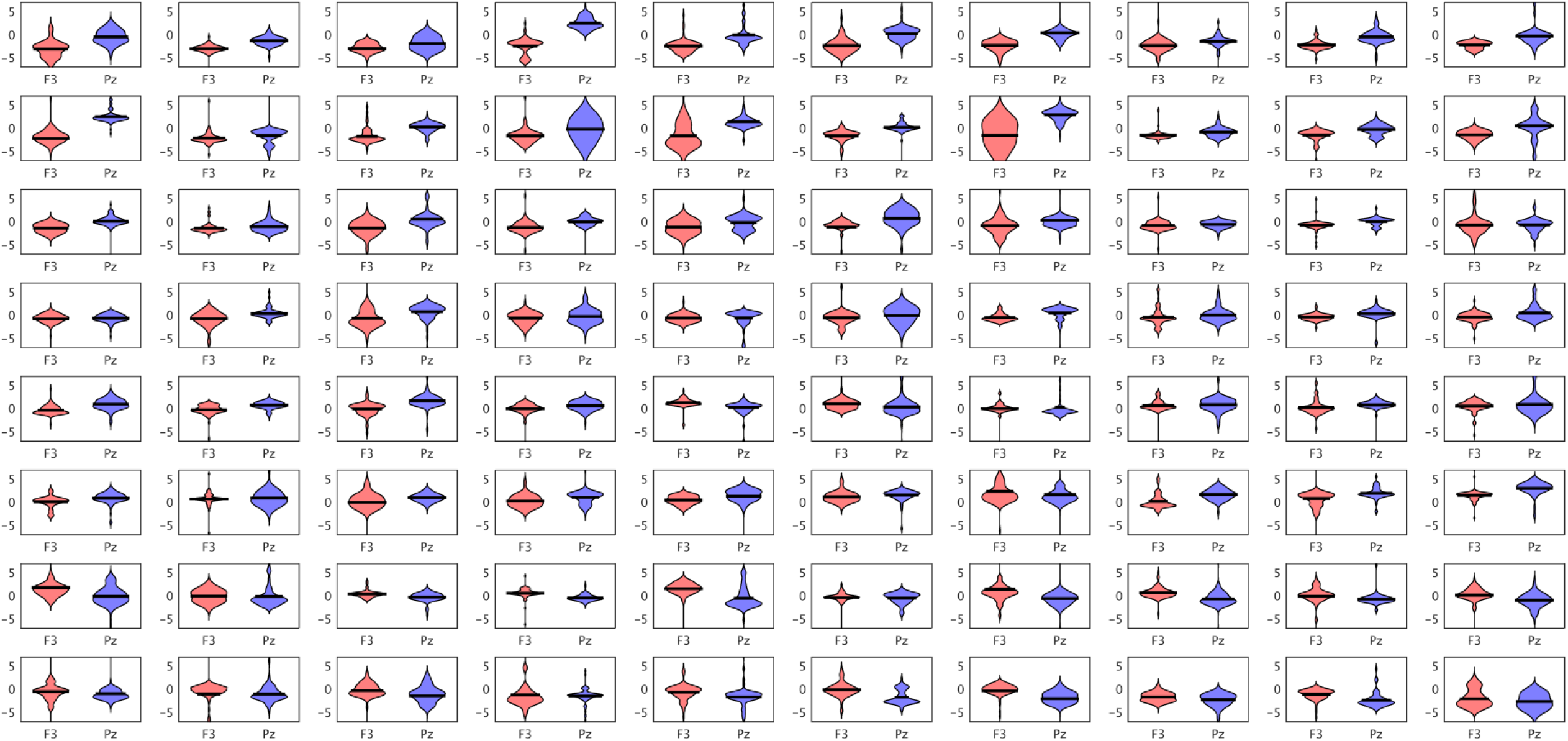

## Supplementary Materials A

The amplitude voltage between 300 and 500 ms recorded at F3 and Pz were extracted for each participant and item. The voltages of each electrode and condition were sorted based on amplitude and divided into 76 bins (as 77 was the minimum number of artifact-free items per condition, 76 bins guaranteed the fewest possible trials per bin, reducing the potential consequences of averaging procedure; each bin contained 1 or 2 trials). When two trials belonged to the same bin they were averaged, and the amplitude voltages of corresponding bins were subtracted across conditions. The resulting ERP effects calculated for F3 and Pz are plotted for each participant as violin plots (black lines represent means) in order to show the distributions of ERP effects calculated on small sets of trials. Y-axis units are microvolts. X-axis shows the electrodes of interest. The subject order is the same as Fig. 2. There is a correspondence between individual topographies shown in Fig. 2 and the distributions of ERP effects at the trial level.

## Supplementary Materials B

Proportions of LAN and N400 effects (300-500 ms) calculated based on different criteria from those described in the paper

### Effects defined on clusters of electrodes

A LAN effect was considered present when the negative effect (disagreement-agreement voltage difference) over a left-anterior cluster of electrodes (F3, F7, FC5) was greater than the effect registered over a posterior cluster (P3, Pz, P4).

|  | Subjects | Items |
| --- | --- | --- |
| LANs | 49% | 44% |
| N400s | 29% | 32% |
| Positivities | 23% | 25% |
| Negative effect at F3, F7, FC5 | 68% | 64% |

A N400 effect was defined as a negative effect over the posterior cluster (P3, Pz, P4) being greater than the effect over the left-anterior cluster (F3, F7, FC5).

Any other case where both left-anterior and posterior cluster showed a positive ERP difference was categorized as positive effect.

### Effects defined by laterality and anteriority indexes

Average ERP effects (disagreement-agreement voltage difference) were calculated for anterior (A: Fp1, Fp2, F3, F4, F7, F8, Fz, FC1, FC2, FC5, FC6), posterior (P: P3, P4, O1, O2, P7, P8, Pz, CP1, CP2, CP5, CP6), left (L: Fp1, F3, C3, P3, O1, F7, T7, P7, FC1, CP1, FC5, CP5) and right electrodes (R: Fp2, F4, C4, P4, O2, F8, T8, P8, FC2, CP2, FC6, CP6).

For each ERP effect a laterality and anteriority index was defined and an index size was quantified based on the following formulas:

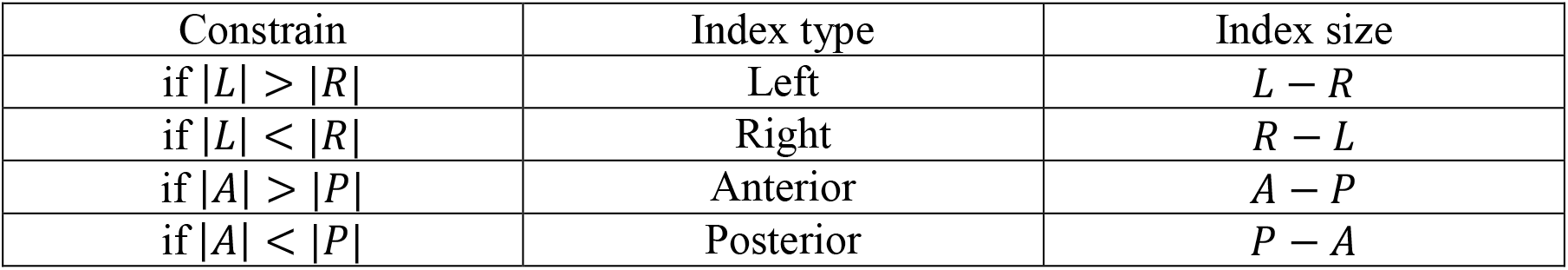

Then, each effect was classified as negative when the sign of laterality and anteriority indexes were negative and positive when the sign of both indexes were positive. Mixed effects (positive and negative) are reported aside.

|  |  | Subjects | Items |
| --- | --- | --- | --- |
| Negative effects | Left anterior (LANs) | 30% | 19% |
|  | Right anterior | 9% | 10% |
|  | Left posterior | 13% | 17% |
|  | Right posterior (N400s) | 6% | 11% |
| Positive effects | Left anterior | 3% | 5% |
|  | Right anterior | 16% | 13% |
|  | Left posterior | 3% | 6% |
|  | Right posterior (P600s) | 15% | 16% |
| Mixed effects | Left neg., Ant. pos. | 1% | 0.5% |
|  | Left pos., Post. neg. | 0% | 1% |
|  | Right pos., Post. neg. | 3% | 0% |
|  | Right neg., Post. pos. | 3% | 0.5% |

## Supplementary Materials C

| LAN (lmer) | Model A |  |  | Model B |  |  | Model C |  |  | Model D |  |  |
| --- | --- | --- | --- | --- | --- | --- | --- | --- | --- | --- | --- | --- |
| | $\beta$ | CI | $t$ | $\beta$ | CI | $t$ | $\beta$ | CI | $t$ | $\beta$ | CI | $t$ |
| **Fixed Parts** |  |  |  |  |  |  |  |  |  |  |  |  |
| Control | 0.04 | [-0.15, 0.23] | 0.44 | 0.05 | [-0.50, 0.59] | 0.17 | 0.05 | [-0.48, 0.57] | 0.17 | 0.05 | [-0.49, 0.58] | 0.17 |
| Disagreement | -0.62 | [-0.87, -0.38] | 4.94 | -0.62 | [-0.86, -0.38] | 5.13 | -0.62 | [-0.88, -0.36] | 4.61 | -0.62 | [-0.89, -0.35] | 4.57 |
| **Random Parts** |  |  |  |  |  |  |  |  |  |  |  |  |
| $\sigma^2$ | | 69.83 | | | 64.40 | | | 64.33 | | | 64.30 | |
| $\tau_{00,i}$ (SD) | | 0.33 (0.58) | | | 0.39 (0.62) | | | 0.39 (0.63) | | | 0.61 (0.78) | |
| $\tau_{00,s}$ (SD) | | | | | 5.41 (2.32) | | | 5.10 (2.26) | | | 5.10 (2.26) | |
| $\tau_{00,dis-s}$ (SD) | | | | | | | | 0.27 (0.52) | | | 0.27 (0.52) | |
| $\tau_{00,dis-i}$ (SD) | | | | | | | | | | | 0.09 (0.30) | |
| $\rho_{01}$ | | | | | | | | | | | | |
| $N_i$ | | 240 | | | 240 | | | 240 | | | 240 | |
| $N_s$ | | | | | 80 | | | 80 | | | 80 | |
| $ICC_i$ | | 0.005 | | | 0.006 | | | 0.006 | | | 0.009 | |
| $ICC_s$ | | | | | 0.077 | | | 0.073 | | | 0.073 | |
| Observations |  | 17523 |  |  | 17523 |  |  | 17523 |  |  | 17523 |  |
| $R^2 / \Omega_0^2$ | | .014 / .010 | | | .092 / .091 | | | .094 / .093 | | | .094 / .093 | |
| **Correlation of Fixed Effects** |  |  |  |  |  |  |  |  |  |  |  |  |
|  |  | -0.65 |  |  | -0.22 |  |  | -0.12 |  |  | -0.14 |  |
| **Model comparison ($\chi^2$)** |  |  |  |  | 1167.41 |  |  | 2.96 |  |  | 2.44 |  |

| LAN (stan) | Model A |  | Model B |  | Model C |  | Model D |  |
| --- | --- | --- | --- | --- | --- | --- | --- | --- |
| | $\beta$ | Cr. I | $\beta$ | Cr. I | $\beta$ | Cr. I | $\beta$ | Cr. I |
| **Fixed Parts** |  |  |  |  |  |  |  |  |
| Control | 0.04 | [-0.15, 0.23] | 0.03 | [-0.51, 0.59] | 0.04 | [-0.48, 0.58] | 0.05 | [-0.49, 0.60] |
| Disagreement | -0.62 | [-0.87, -0.38] | -0.62 | [-0.86, -0.39] | -0.62 | [-0.87, -0.35] | -0.62 | [-0.89, -0.34] |
| **Random Parts** |  |  |  |  |  |  |  |  |
| $\sigma^2$ | | 69.84 | | 64.43 | | 64.35 | | 64.34 |
| SD $\tau_{00,i}$ | | 0.58 | | 0.62 | | 0.62 | | 0.69 |
| SD $\tau_{00,s}$ | | | | 2.37 | | 2.29 | | 2.29 |
| SD $\tau_{00,dis-s}$ | | | | | | 0.56 | | 0.57 |
| SD $\tau_{00,dis-i}$ | | | | | | | | 0.35 |
| $\rho_{01}$ | | | | | | | | |
| $N_i$ | | 240 | | 240 | | 240 | | 240 |
| $N_s$ | | | | 80 | | 80 | | 80 |
| Observations |  | 17523 |  | 17523 |  | 17523 |  | 17523 |
| $R^2$ | | 0.006 | | 0.083 | | 0.084 | | 0.085 |
| **Model comparison (elpd, SE)** |  |  |  | 666.3, 37.8 |  | -0.6, 2.3 |  | 0.6, 0.9 |

Note: CI, confidence interval; Cr. I, credibility interval.

Fixed and random effect parameters are reported for lmer models (upper side) and Stan models (lower side) of LAN variability. Model comparisons were carried out using Chi-squared test in the case of lmer models and using leave-one-out cross validation in the case of Stan models (this method does not have the mathematical limitations of the likelihood ratio test, as pointed out in Bates, 2010; Demidenko, 2013; Pinheiro & Bates, 2000). The fixed effect parameter estimates are similar in the two approaches. Conclusions for the model comparisons based on either the like-hood ratio test or the leave-one-out cross validation are the same.

Here below fixed and random effect parameters are reported for lmer models of P600 variability.

| **P600 (lmer)** | **Model A** |  |  | **Model B** |  |  | **Model C** |  |  | **Model D\*** |  |  |
| --- | --- | --- | --- | --- | --- | --- | --- | --- | --- | --- | --- | --- |
| | $\beta$ | CI | $t$ | $\beta$ | CI | $t$ | $\beta$ | CI | $t$ | $\beta$ | CI | $t$ |
| **Fixed Parts** |  |  |  |  |  |  |  |  |  |  |  |  |
| Control | 0.81 | [0.59, 1.02] | 7.40 | 0.79 | [0.31, 1.28] | 3.24 | 0.81 | [0.43, 1.18] | 4.24 | 0.81 | [0.45, 1.17] | 4.44 |
| Disagreement | 3.69 | [3.42, 3.96] | 26.89 | 3.70 | [3.43, 3.96] | 27.59 | 3.68 | [3.13, 4.22] | 13.20 | 3.67 | [3.10, 4.24] | 12.71 |
| **Random Parts** |  |  |  |  |  |  |  |  |  |  |  |  |
| $\sigma^2$ | | 82.32 | | | 78.48 | | | 77.282 | | | 77.276 | |
| $\tau_{00,i}$ (SD) | | 0.62 (0.79) | | | 0.67 (0.82) | | | 0.70 (0.84) | | | 0.00 (0.00) | |
| $\tau_{00,s}$ (SD) | | | | | 3.87 (1.97) | | | 1.95 (1.40) | | | 1.95 (1.40) | |
| $\tau_{00,dis-s}$ (SD) | | | | | | | | 4.78 (2.19) | | | 4.78 (2.19) | |
| $\tau_{00,dis-i}$ (SD) | | | | | | | | | | | 1.42 (1.19) | |
| $\rho_{01}$ | | | | | | | | | | | | |
| $N_i$ | | 240 | | | 240 | | | 240 | | | 240 | |
| $N_s$ | | | | | 80 | | | 80 | | | 80 | |
| $ICC_i$ | | 0.008 | | | 0.008 | | | 0.009 | | | 0.000 | |
| $ICC_s$ | | | | | 0.047 | | | 0.024 | | | 0.025 | |
| Observations |  | 17523 |  |  | 17523 |  |  | 17523 |  |  | 17523 |  |
| $R^2 / \Omega_0^2$ | | .052 / .051 | | | .100 / .099 | | | .117 / .116 | | | .117 / .116 | |
| **Correlation of Fixed Effects** |  |  |  |  |  |  |  |  |  |  |  |  |
|  |  | -0.62 |  |  | -0.27 |  |  | 0.02 |  |  | 0.02 |  |
| **Model comparison ($\chi^2$)** |  |  |  |  | 629.55 |  |  | 182.72 |  |  | 0.81 |  |

Variance-Covariance matrices

**LAN**

|  | Control | Disagreement |
| --- | --- | --- |
| Control | 0.0093 | -0.0079 |
| Disagreement | -0.0079 | 0.0160 |

Model B<-lmer(vLAN ~ agr + (1 | s) + (1 | i),data=d)
|  | Control | Disagreement |
| --- | --- | --- |
| Control | 0.0765 | -0.0073 |
| Disagreement | -0.0073 | 0.0147 |

Model C<-lmer(vLAN ~ agr + (1 + agr | s) + (1 | i),data=d)
|  | Control | Disagreement |
| --- | --- | --- |
| Control | 0.0727 | -0.0043 |
| Disagreement | -0.0043 | 0.0181 |

Model D<-lmer(vLAN ~ agr + (1 + agr | s) + (1 + agr | i),data=d)
|  | Control | Disagreement |
| --- | --- | --- |
| Control | 0.0735 | -0.0052 |
| Disagreement | -0.0052 | 0.0184 |

**P600**

P600 Model A<-lmer(vP600 ~ agr + (1 | i),data=d)
|  | Control | Disagreement |
| --- | --- | --- |
| Control | 0.0119 | -0.0093 |
| Disagreement | -0.0093 | 0.0188 |

Model B<-lmer(vP600 ~ agr + (1 | s) + (1 | i),data=d)
|  | Control | Disagreement |
| --- | --- | --- |
| Control | 0.0600 | -0.0089 |
| Disagreement | -0.0089 | 0.0179 |

Model C<-lmer(vP600 ~ agr + (1 + agr | s) + (1 | i),data=d)
|  | Control | Disagreement |
| --- | --- | --- |
| Control | 0.0361 | 0.0008 |
| Disagreement | 0.0008 | 0.0775 |

Model D<-lmer(vP600 ~ agr + (1 + agr | s) + (1 + agr | i),data=d)\*
|  | Control | Disagreement |
| --- | --- | --- |
| Control | 0.0332 | 0.0010 |
| Disagreement | 0.0010 | 0.0834 |

\*model did not converge

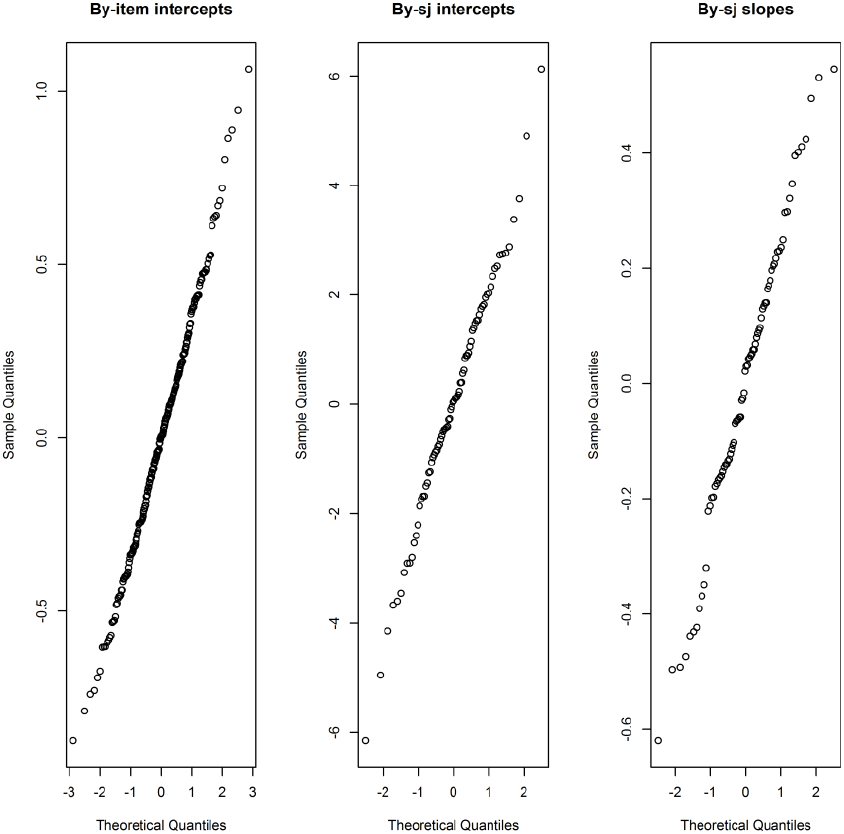

## Supplementary Materials D

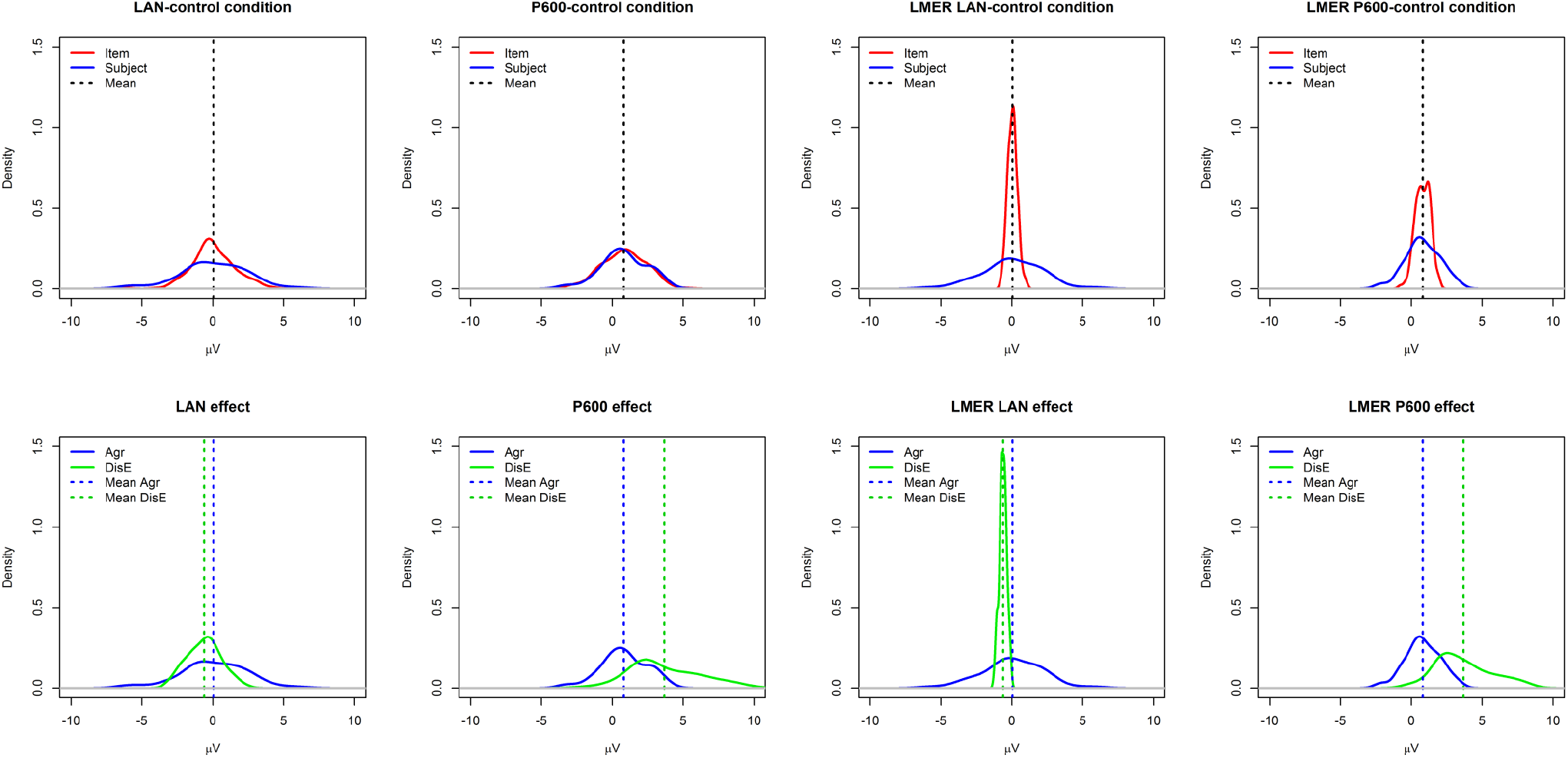

Density plots calculated on the real data (four graphs on the left) and on the variance parameters estimated from models C (four graphs on the right). Y-axis are kernel density estimates, and x-axis units are microvolts. The curves represent the estimates of the population probability distribution based on the given sample of points.

The upper panel shows the LAN and the P600 average response to the control condition (vertical dashed line), and variability due to subject (blue distribution) and item (red distribution). The lower panel shows the ERP average response to the control condition (Mean Agr: blue dashed line) and the ERP average size of the violation effect (Mean DisE: green dashed line) with the relative variability due to subjects.

Note that the distributions based on the estimates from the mixed-linear models show less variability as compared to those based on the real data. This is due to the different sources of variance represented here. The observed data distributions do not distinguish the sources of random effects. Thus, the variance of these distributions can include measurement variability (e.g., variations in the underlying response from subject to subject) as well as measurement noise. The mixed-linear model distributions showed only the measurement variability specifically related to items or subjects variations. Mixed-linear models have the advantage to tease apart measurement variability and measurement noise, showing the magnitude of the random effects specifically associated with individual differences.

## References

Baayen, R. H., Davidson, D. J., & Bates, D. M. (2008). Mixed-effects modeling with crossed random effects for subjects and items. Journal of memory and language, 59(4), 390–412. doi: 10.1016/j.jml.2007.12.005

Barr, D.J., Levy, R., Scheepers, C., & Tily, H.J. (2013). Random effects structure for confirmatory hypothesis testing: Keep it maximal. Journal of Memory and Language, 68, 255–278. doi: 10.1016/j.jml.2012.11.001

Bates, D. M., Mächler, M., Bolker, B. M., & Walker, S. C. (2014). lme4: Linear Mixed-Effects Models Using Eigen and S4. R Package Version 1.1-7. Available online at: http://CRAN.R-project.org/package=lme4

Caffarra, S., & Barber, H. (2015). Does the ending matter? The role of gender-to-ending consistency in sentence reading. Brain Research, 1605, 83–92. doi: 10.1016/j.brainres.2015.02.018

Caffarra, S., Molinaro, N., Davidson, D., & Carreiras, M. (2015). Second language syntactic processing revealed through event-related potentials: An empirical review. Neuroscience & Biobehavioral reviews, 51C, 31–47. doi:10.1016/j.neubiorev.2015.01.010

Caffarra, S., Barber, H., Molinaro, N., & Carreiras, M. (2017). When the end matters: influence of gender cues during agreement computation in bilinguals. Language, Cognition and Neuroscience. doi: 10.1080/23273798.2017.1283426

Cooper, R., Osselton, J. W., & Shaw, J. C. (1969). EEG technology. London, UK: Butterworths.

Coulson, S., King, J.W., & Kutas, M. (1998). Expect the Unexpected: Event-related Brain Response to Morphosyntactic Violations. Language and cognitive processes, 1998, 13(1), 21–58. doi: 10.1080/016909698386582

Friederici, A. D. (2002). Towards a neural basis of auditory sentence processing. Trend in Cognitive Sciences, 6 (2), 78–84. doi: 10.1016/S1364-6613(00)01839-8

Friston, K. (2012). The ironic rules for non-statistical reviewers. NeuroImage, 61, 1300–1310. doi: 10.1016/j.neuroimage.2012.04.018

Gelman, A., & Carlin, J. (2014). Beyond Power Calculations: Assessing Type S (Sign) and Type M (Magnitude) Errors. Perspectives on Psychological Science, 9, 641–651. doi: 10.1177/1745691614551642

Gunter, T. C., Friederici, A. D., & Schriefers, H. (2000). Syntactic gender and semantic expectancy: ERPs reveal early autonomy and late interaction. Journal of Cognitive Neuroscience, 12, 556–568. doi: 10.1162/089892900562336

Hagoort, P. (2003). Interplay between syntax and semantics during sentence comprehension: ERP effects of combining syntactic and semantic violations. Journal of Cognitive Neuroscience, 15, 883–899. doi:10.1162/089892903322370807

Hallez, H., Vanrumste, B., Grech, R., Muscat, J., De Clercq, W., Vergult, A. … Lemahieu, I. (2007). Review on solving the forward problem in EEG source analysis. Journal of NeuroEngineering and Rehabilitation, 4, 107–111. doi: 10.1186/1743-0003-4-46

Kemmerer, D. (2015). Cognitive Neuroscience of Language. New York: Psychology Press.

Krott, A., & Lebib, R. (2013). Electrophysiological evidence for a neural substrate of morphological rule application in correct wordforms, Brain Research, 1496, 70–83. doi: 10.1016/j.brainres.2012.12.012

Kutas, M., & Federmeier, K. D. (2011). Thirty years and counting: finding meaning in the N400 component of the event-related brain potential (ERP). Annual Review of Psychology, 62, 621–647. doi: 10.1146/annurev.psych.093008.131123.

Lindquist, M. A., Caffo, B., & Crainiceanu, C. (2013). Ironing out the statistical wrinkles in “Ten Ironic Rules”. NeuroImage, 81, 499–502. doi: 10.1016/j.neuroimage.2013.02.056

Loken, E., & Gelman, A. (2017). Measurement error and replication crisis. Science, 355, 584–585. doi: 10.1126/science.aal3618

Luck, S. J. (2005). An introduction to the Event-Related Potential Technique. Cambridge, MA: MIT Press.

Mendoza, M. (2017). Morphosyntactic processing at the subject-level: ERP evidence on the left anterior negativity (LAN) effect. BCBL Master thesis 2016-2017.

Molinaro, N., Barber, H., & Carreiras, M. (2011). Grammatical agreement processing in reading: ERP findings and future directions. Cortex, 47, 908–930. doi: 10.1016/j.cortex.2011.02.019

Molinaro, N., Barber, H.A., Caffarra, S., & Carreiras, M. (2015). On the left anterior negativity (LAN): The case of morphosyntactic agreement. Cortex, 66, 156–159. doi: 10.1016/j.cortex.2014.06.009

Osterhout, L. (1997).On the brain response to syntactic anomalies: manipulations of word position and word class reveal individual differences. Brain & Language, 59, 494–522. doi: 10.1006/brln.1997.1793

Osterhout, L., & Mobley, L. (1995). Event-related brain potentials elicited by failure to agree. Journal of Memory and Language, 34, 739–773. doi: 10.1006/jmla.1995.1033

Osterhout, L., & Hagoort, P. (1999). A superficial resemblance does not necessarily mean you are part of the family: Counterarguments to Coulson, King and Kutas (1998) in the P600/SPS-P300 debate. Language and Cognitive Processes, 14, 1–14. doi: 10.1080/016909699386356

Osterhout, L., McLaughlin, J., Kim, A., Greenwald, R., & Inoue, K. (2004). Sentences in the brain: Event-related potentials as real-time reflections of sentence comprehension and language learning. In M. Carreiras & C. Clifton (Eds.), The On-line Study of Sentence Comprehension: Eyetracking, ERPs and Beyond. (pp. 271–308). New York: Psychology Press. doi: 10.4324/9780203509050

Pinheiro, J. C., & Bates, D. M. (2000). Linear mixed-effects models: basic concepts and examples. Mixed-effects models in S and S-Plus, 3–56. New York: Springer.

Roll, M., Gosselke, S., Lindgren, M., & Horne, M. (2013). Time-driven effects on processing grammatical agreement. Frontiers in Psychology, 4, 1–8. doi: 10.3389/fpsyg.2013.01004

Sörnmo, L., & Laguna, P. (2005). Bioelectrical signal processing in cardiac and neurological applications. San Diego, US: Elsevier Academic Press.

Tanner, D. (2015). On the left anterior negativity (LAN) in electrophysiological studies of morphosyntactic agreement: a commentary on “grammatical agreement processing in reading: ERP findings and future directions” by Molinaro et al., 2014. Cortex, 66, 149–155. doi: 10.1016/j.cortex.2014.04.007

Tanner, D., McLaughlin, J., Herschensohn, J., & Osterhout, L. (2013). Individual differences reveal stages of L2 grammatical acquisition: ERP Evidence. Bilingualism: Language and Cognition, 16(2), 367–382. doi: 10.1017/S1366728912000302

Tanner, D., Inoue, K., & Osterhout, L. (2014). Brain-based Individual Differences in On-line L2 Grammatical Comprehension. Bilingualism: Language and Cognition, 17(2), 277–293. doi: 10.1017/S1366728913000370

Tanner, D., & Van Hell, J. G. (2014). ERPs reveal individual differences in morphosyntactic processing. Neuropsychologia, 56, 289–301. doi:10.1016/j.neuropsychologia.2014.02.002

Wicha, N. Y. Y., Moreno, E. M., & Kutas, M. (2004). Anticipating words and their gender: An event-related brain potential study of semantic integration, gender expectancy, and gender agreement in Spanish sentence reading. Journal of Cognitive Neuroscience, 16, 1272–88. doi: 10.1162/0898929041920487

